# Canonical EEG Microstate Dynamic Properties and Their Associations with fMRI Signals at Resting Brain

**DOI:** 10.1101/2020.08.14.251066

**Authors:** Obada Al Zoubi, Masaya Misaki, Aki Tsuchiyagaito, Ahmad Mayeli, Vadim Zotev, Tulsa 1000 Investigators, Hazem Refai, Martin Paulus, Jerzy Bodurka

**Affiliations:** Laureate Institute for Brain Research, Tulsa, OK, United States; Japan Society for the Promotion of Science, Tokyo, Japan; Department of Community Medicine, Oxley Health Sciences, University of Tulsa, Tulsa, Oklahoma, United States; Stephenson School of Biomedical Engineering, University of Oklahoma, Norman, OK, United States; Electrical and Computer Engineering, University of Oklahoma, Tulsa, OK, United States

## Abstract

Electroencephalography microstates (EEG-ms) capture and reflect the spatio-temporal neural dynamics of the brain. A growing literature is employing EEG-ms-based analyses to study various mental illnesses and to evaluate brain mechanisms implicated in cognitive and emotional processing. The spatial and functional interpretation of the EEG-ms is still being investigated. Previous works studied the association of EEG-ms time courses with blood-oxygen-level-dependent (BOLD) functional magnetic resonance imaging (fMRI) signal and suggested an association between EEG-ms and resting-state networks (RSNs). However, the distinctive association between EEG-ms temporal dynamics and brain neuronal activities is still not clear, despite the assumption that EEG-ms are an electrophysiological representation of RSNs activity. Recent works suggest a role for brain spontaneous EEG rhythms in contributing to and modulating canonical EEG-ms topographies and determining their classes (coined A through D) and metrics. This work simultaneously utilized EEG and fMRI to understand the EEG-ms and their properties further. We adopted the canonical EEG-ms analysis to extract three types of regressors for EEG-informed fMRI analyses: EEG-ms direct time courses, temporal activity per microstate, and pairwise temporal transitions among microstates (the latter two coined activity regressors). After convolving EEG-ms regressors with a hemodynamic response function, a generalized linear model whole-brain voxel-wise analysis was conducted to associate EEG-ms regressors with fMRI signals. The direct time course regressors replicated prior findings of the association between the fMRI signal and EEG-ms time courses but to a smaller extent. Notably, EEG-ms activity regressors were mostly anticorrelated with fMRI, including brain regions in the somatomotor, visual, dorsal attention, and ventral attention fMRI networks with no significant overlap for default mode, limbic or frontoparietal networks. A similar pattern emerged in using the transition regressors among microstates but not in self-transitions. The relatively short duration of each EEG-ms and the significant association of EEG-ms activity regressors with fMRI signals suggest that EEG-ms manifests successive transition from one brain functional state to another rather than being associated with specific brain functional state or RSN networks.

## Introduction

Electroencephalography (EEG) is a direct measure of brain neuronal activity with a high temporal resolution (e.g., order of milliseconds), although it suffers from low spatial resolution. Distinct topographic representations of the EEG electric scalp potentials—lasting a few dozens of a millisecond, termed EEG-microstates (EEG-ms)—provide a novel opportunity for the search of potential biomarkers of various psychiatric disorders (Khanna, Pascual-Leone, Michel, & Farzan, 2015). Understanding the spatial and functional brain representation of EEG-ms is an active field of research. One accepted notion is that EEG-ms may represent the coordinated neuronal current activity in space and time (Khanna et al., 2015; Michel & Koenig, 2018). Thus, a change in the topographies of EEG-ms may be attributed to a change in the orientation or distribution of the current dipoles underlying EEG signal formation (Dr Lehmann, Ozaki, & Pal, 1987; Vaughan Jr, 1980). In addition, EEG-ms are correlated with several mental processes including attention, language processing, perceptual awareness, various resting-state networks and visual processing (Brandeis & Lehmann, 1989; Brandeis, Lehmann, Michel, & Mingrone, 1995; Britz, Díaz Hernàndez, Ro, & Michel, 2014; Britz & Michel, 2010; Britz, Van De Ville, & Michel, 2010; Koenig & Lehmann, 1996; Michel & Koenig, 2018; Michel et al., 2001; Pizzagalli, Lehmann, König, Regard, & Pascual-Marqui, 2000). Furthermore, EEG-ms has been used to characterize different mental disorders such as schizophrenia, Alzheimer’s disease, dementia, mood\anxiety disorders, major depressive disorders, and autism (Al Zoubi et al., 2019; Andreou et al., 2014; da Cruz et al., 2020; Damborská et al., 2019; Dierks et al., 1997; Drissi et al., 2016; D’Croz-Baron, Baker, Michel, & Karp, 2019; Jia & Yu, 2019; Kikuchi et al., 2011; Dietrich Lehmann et al., 2005; Murphy et al., 2020; Nishida et al., 2013; Stevens & Kircher, 1998; W. Strik, Dierks, Becker, & Lehmann, 1995; W. K. Strik et al., 1997; Tomescu et al., 2014). EEG-ms has high temporal resolution; however, it suffers from low spatial resolution. Thus, several studies have investigated the association between EEG-ms (Dr Lehmann et al., 1987; Yuan et al., 2018; Yuan, Zotev, Phillips, Drevets, & Bodurka, 2012) and functional magnetic resonance imaging (fMRI) signals, revealing a correlation between the EEG-ms time course and fMRI resting-state networks (RSNs) (Britz et al., 2010; Musso, Brinkmeyer, Mobascher, Warbrick, & Winterer, 2010; Yuan et al., 2012). Moreover, the source localization of EEG-ms was investigated in (Custo et al., 2017), where authors identified seven EEG-ms (A through G) and localized their sources from healthy volunteers. Their results suggested a common fMRI activation among EEG-ms in several of the brain’s hubs, e.g., anterior and posterior cingulate cortices, insula, superior frontal cortex, and precuneus. Other works have used the voxel-wise generalized linear model (GLM) analysis to associate EEG-ms time courses with blood-oxygen-level-dependent (BOLD) signals. For instance, using canonical EEG-ms analysis (Michel & Koenig, 2018), the authors in (Britz et al., 2010) demonstrated that for resting-state fMRI: 1) EEG-ms A (MS-A) is negatively correlated with the BOLD signal in the bilateral superior and middle temporal lobe; 2) EEG-ms B (MS-B) is negatively correlated with the BOLD signal in the bilateral occipital cortex; 3) EEG-ms C (MS-C) is positively correlated with the BOLD signal in the right insular cortex, bilateral inferior frontal cortices and the dorsal anterior cingulate cortex; and 4) EEG-ms D (MS-D) is negatively associated with BOLD signal within frontoparietal regions. Similarly, authors (Musso et al., 2010) extracted 10 EEG-ms from healthy volunteers, revealing a significant association between the spatial maps of EEG-ms and BOLD signals in multiple brain regions. Using frequency-varying EEG-ms information, the authors in (Schwab et al., 2015) found that thalamic nuclei subregions exhibit different BOLD fMRI activation patterns based on the employed frequency.

The previous EEG-fMRI results suggested that EEG-ms are associated with one or more brain RSNs activities, and the brain network activities sum up and contribute to a specific EEG-ms class. However, this assumption was recently reconsidered by studying the impact of the intracortical sources of EEG alpha oscillation on the topography of the EEG-ms classes (Milz, Pascual-Marqui, Achermann, Kochi, & Faber, 2017). The results revealed that the intracortical strength of alpha oscillation and its distribution predominantly determined the EEG-ms topographies. Additionally, the authors in (Croce, Quercia, Costa, & Zappasodi, 2020) showed that EEG-ms occurrence, coverage, and duration are influenced by EEG alpha oscillation, and are sensitive to intra- and inter-subject alpha rhythm variability, further indicating a prominent influence of brain oscillation on the EEG-ms. The alpha rhythm was shown to decrease during task-related mental activity (Goldman, Stern, Engel Jr, & Cohen, 2002; G Pfurtscheller, 2003; Salenius, Kajola, Thompson, Kosslyn, & Hari, 1995; Scheeringa, Petersson, Kleinschmidt, Jensen, & Bastiaansen, 2012). Additionally, alpha rhythm has been shown to have a modularity effect on the BOLD signal (Mantini, Perrucci, Del Gratta, Romani, & Corbetta, 2007; Scheeringa et al., 2012) while exhibiting an anticorrelation relationship epically for the occipital cortex, observed in fMRI and other imaging modalities (Goldman et al., 2002; Jacquy, Charles, Piraux, & Noel, 1980; H Laufs et al., 2003; Moosmann et al., 2003; Sadato et al., 1998). That is, we posit that the interaction among BOLD signals, EEG alpha rhythm, and EEG-ms may suggest that EEG-ms represents transitions among brain states or network activities rather than representation of the specific brain states or network activities.

We therefore investigated the association between EEG-ms dynamics and whole brain voxel-wise BOLD signals to assess the assumption that EEG-ms represent specific a brain function or network. To do so, we extracted the time course of the four canonical EEG-ms and used them as regressors in the GLM. This allowed us to evaluate the spatial distribution of the canonical EEG-ms in the fMRI domain. Furthermore, we evaluated whether EEG-ms are associated with transition among different brain functions or network activities by introducing two new sets of regressors that categorize the temporal dynamics of EEG-ms: 16 pairwise transition probabilities among EEG-ms (including four self-transition) and four regressors of switching activity per each EEG-ms.

## Methods

### Participants

We selected 52 healthy control subjects (mean age: 32, standard deviation (SD): 12 years; 25 females) from the first 500 subjects of the Tulsa 1000 (T-1000), a naturalistic study assessing and longitudinally following 1000 individuals, including healthy individuals and treatment-seeking individuals with substance use, eating, mood, and/or anxiety disorders (Victor et al., 2018). An 8-min simultaneous EEG-fMRI recording was obtained from each subject during a resting-state scan. The participants were asked to relax and keep their eyes open and fixate their eyes on a cross displayed on the fMRI stimulus projection screen. Further quality control for the simultaneous EEG-fMRI data (see below) reduced the number of subjects to 40 participants (age mean ±SD: 32 ± 12 years; 23 females).

### Simultaneous EEG-fMRI Data Acquisition

MRI imaging and simultaneous EEG-fMRI was conducted on a General Electric Discovery MR750 whole-body 3T MRI scanner with a standard 8-channel, receive-only head coil array. A single-shot gradient-recalled echoplanar imaging (EPI) sequence with Sensitivity Encoding (SENSE) was acquired with the following parameters: TR = 2000 ms, TE = 27 ms, FA = 40°, FOV = 240 mm, 37 axial slices with 2.9 mm thickness with 0.5 mm gap, matrix = 96×96, SENSE acceleration factor R = 2. EPI images were reconstructed into a 128×128 matrix that produced 1.875×1.875×2.9 mm3 voxel volume. The resting fMRI run time was 8 min 8 s (244 volumes). To provide anatomical reference, T1-weighted MRI images were acquired using a magnetization-prepared rapid gradient-echo (MPRAGE) sequence with the following parameters: FOV = 240×192 mm, matrix = 256×256, 120 axial slices, slice thickness= 1.1 mm, 0.938×0.938×1.1 mm3 voxel volume, TR = 5 ms, TE = 1.948 ms, R = 2, flip angle =8°, delay time = 1400 ms, inversion time = 725 ms, sampling bandwidth = 31.2 kHz, scan time = 5 min 4 s. During each scan, a pneumatic belt placed around the subject’s torso was used to record respiration, and a photoplethysmograph with an infrared emitter placed under the pad of the subject’s index finger was used to record pulse oximetry. The resting-state fMRI scan was the first scan of each session. During each resting-state scan, subjects were instructed to remain still with their eyes open while looking at a fixation cross on the screen. The only other instruction provided was to “try to clear your mind and don’t think about anything in particular.”

EEG signals were recorded simultaneously with fMRI using a 32-channel, MR-compatible EEG system (Brain Products GmbH) with electrodes arranged according to the international 10–20 system. Electrocardiogram (ECG) signal was recorded using an electrode on the subject’s back. In order to synchronize the EEG system clock with the 10 MHz MRI scanner clock, a Brain Products’ SyncBox device was utilized. EEG temporal and measurement resolutions were 0.2 ms (i.e., 16-bit 5 kS/s sampling) and 0.1 μV, respectively. Hardware filtering throughout acquisition in a frequency band between 0.016 and 250 Hz was applied to EEG signals.

### EEG Data Preprocessing

The following preprocessing steps were performed in BrainVision Analyzer 2 software, as described in (Mayeli, Zotev, Refai, & Bodurka, 2016). More specifically, MRI imaging artifacts within the EEG signal were reduced using the average artifact subtraction (AAS) method (Allen, Josephs, & Turner, 2000). After downsampling EEG signals to 250 Hz, band-rejection filters with 1Hz bandwidth were applied to reduce fMRI slice selection fundamental frequency (19.5 Hz) and its harmonics, mechanical vibration noise (26 Hz) along with an alternating current power line noise (60 Hz). Another bandpass filter from 0.1 to 80 Hz (48 dB/octave) was applied to the EEG data. Ballistocardiogram (BCG) artifacts also were reduced using AAS (Allen, Polizzi, Krakow, Fish, & Lemieux, 1998). EEG was then decomposed using the independent component analysis (ICA) Infomax algorithm (Bell & Sejnowski, 1995) implemented in Analyzer. Power spectrum density, topographic maps, time course signals, kurtosis values, and energy values were analyzed for detecting and removing artifactual ICs, including imaging and residual BCG, muscle, and ocular artifacts. Finally, a back-projection (i.e., inverse ICA) was applied after selecting ICs related to neural activities.

### EEG-ms analysis

The standard spatially independent EEG-ms analysis described in (Michel & Koenig, 2018) was conducted. EEG was referenced using an average-reference (Michel & Koenig, 2018). The number of desired EEG-ms was set to k = 4. The following steps were required before running the clustering algorithm: first, the global field power (GFP) for each subject was calculated from band-passed filtered EEG data between 2 and 20 Hz (or 1 and 40 Hz, based on the experiment). GFP peaks were then identified after smoothing the data with a Gaussian-weighted moving average of 5 samples. Finally, to offer a higher level of accuracy, we randomly selected up to n = 10000 peaks and extracted the corresponding EEG points for later analysis. The selected EEG points were then submitted to the Atomize and Agglomerate Hierarchical Clustering (AAHC) algorithm (Maimon & Rokach, 2005; Murray, Brunet, & Michel, 2008) to identify EEG-ms with k = 4. Next, the group means of EEG-ms were computed by first sorting individual EEG-ms and then finding the common topography across all subjects.

### Extracting EEG-ms Regressor

First, we revisit EEG-ms modeling and extraction to elaborate on the extracted EEG-ms regressors. Canonical EEG-ms analysis relies on segmenting EEG points according to their spatial similarities. Although EEG points can be selected randomly, GFP is commonly used to select potential EEG points for segmentation. GFP is defined as the spatial standard deviation of EEG signals across all channels. It has been shown that the peaks of GFP maintain a high signal-to-noise ratio (Khanna et al., 2015; Dr Lehmann et al., 1987). Thus, focusing on GFP peaks to select potential EEG points for segmentation may improve results (see Figure 1). Furthermore, taking GFP peaks into consideration helps reduce the dimensionality of EEG data.

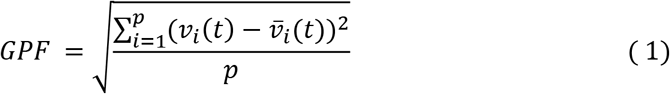

where *p* is the number of electrodes; 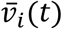 is the mean of electrode values at time point t; and *v_i_*(*t*) is the values of electrode i at time point t.

**Figure 1.**
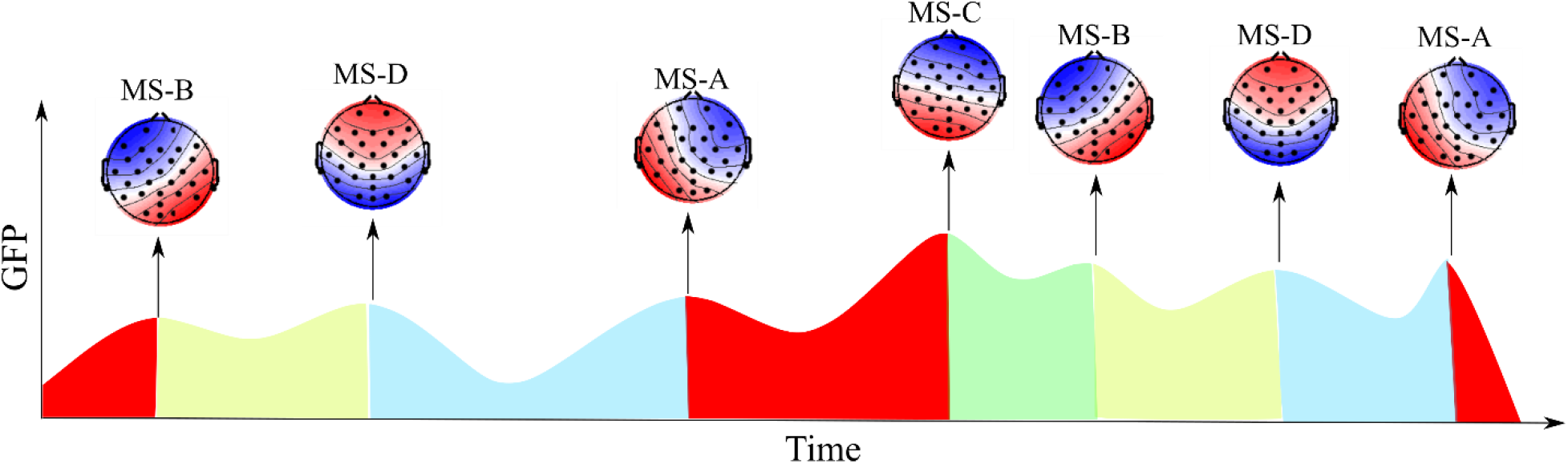
EEG-ms segmentation over time. During any recording of EEG, each point can be assigned to one of four canonical EEG-ms classes.

In this work, we constructed and employed three types of regressors for EEG-informed fMRI analysis: 1) A through D EEG-ms direct time courses; 2) activity per microstate, which measures the extent of switching out of each EEG-ms; and 3) the transition activity between each pair of EEG-ms. It should be noted that the term *time course* here implies a different meaning from other methods that involve time course extraction, such as ICA. EEG-ms time course reflects the spatial similarity between each EEG-ms template (MS-A, MS-B, MS-C, and MS-D) and topographical representation of EEG points. Another difference that arises with the definition of the EEG-ms time course is the polarity consideration of EEG-ms, wherein different interpretations can be drawn if polarity was to be considered.

To provide a better understanding of the time course of EEG-ms, the following section describes the mathematical representation of EEG-ms regressors. First, let’s consider ***x_t_*** to be the electrodes’ value at time *t*. EEG-ms analysis assumes that each EEG point can be presented as:

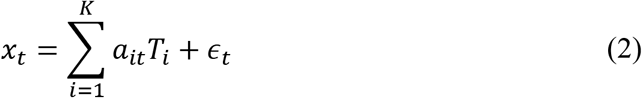

where ***x_t_*** are the electrodes’ value vector at time *t*; *a_it_* is a factor related to each EEG-ms at each time point; and *ϵ_t_* is the error term associated with assigning that time point to one particular EEG-ms (i.e., noise due to the lack of explained topographical representation of that point by the assigned MSs template *K* is the number of the assumed EEG-ms patterns). *T_i_* is the template of EEG-ms *i*.

The time course of EEG-ms is the similarity among each EEG point and each specific EEG-ms template, and can be given as follows:

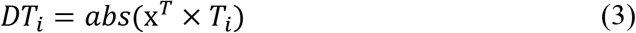

The absolute term in the equation accounts for the polarity invariant property of EEG-ms analysis. **Fig. 2** shows an example of the resulting regressors.

**Figure 2.**
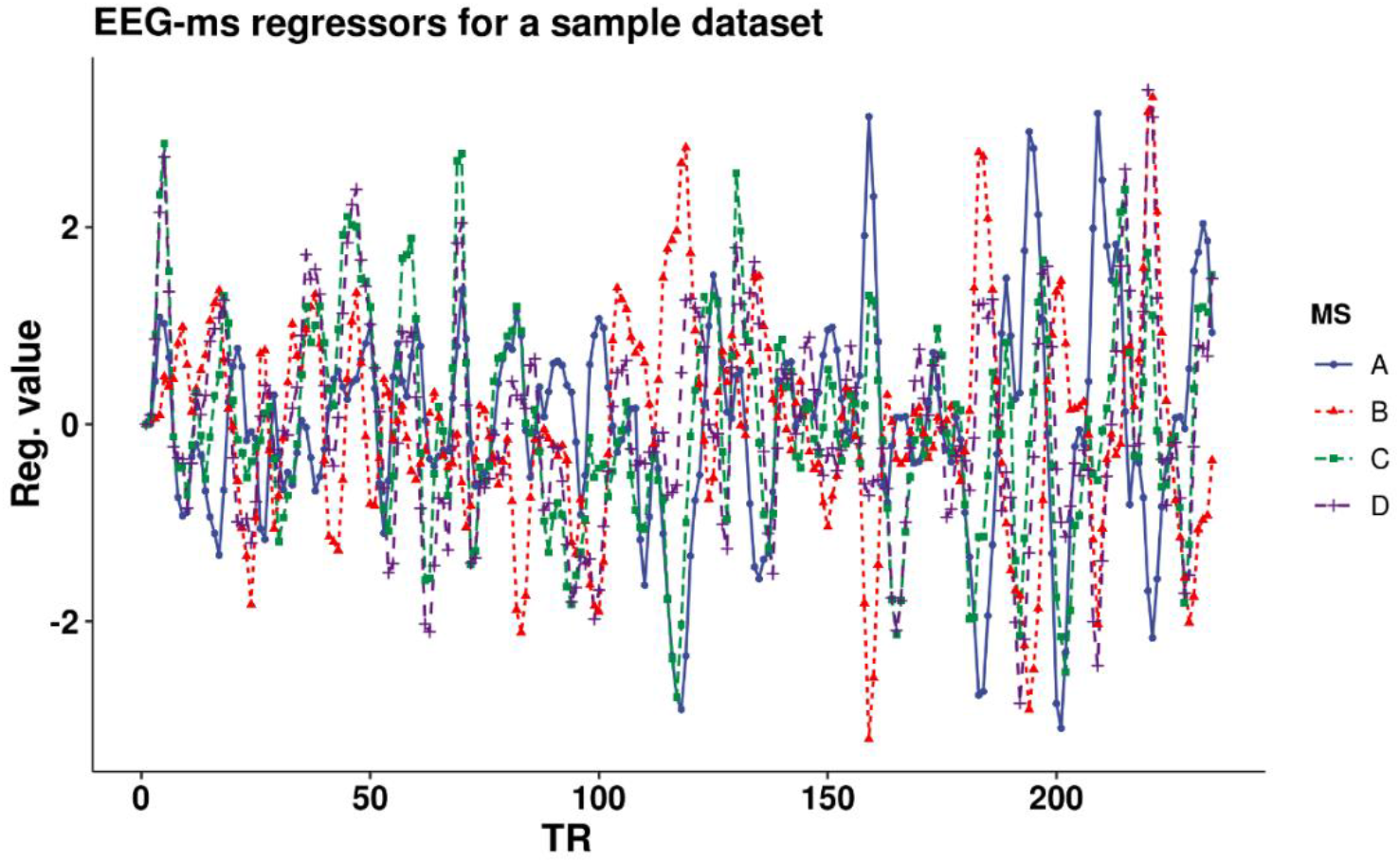
An example of A through D EEG-ms direct time course regressors.

In addition to the direct time course regressors, the activity of each EEG-ms was extracted using a sliding window. The activity metric measures the switches over time from one microstate to other ones and further reveals the brain regions active during the switching of each microstate. Additionally, we investigated the association between switching among pairs of microstates and BOLD level signal change. Mathematically, first, we label EEG points using winner-take-all, i.e., EEG points are assigned to the microstate with the highest similarity as follows:

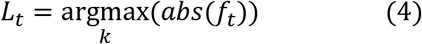

*L_i_* is the assigned label for each EEG point using a winner-takes-all approach, and *i* is the EEG point index. The EEG-ms activity regressors are calculated as the summation of the number of switches from microstate *k* to other microstates in a window size of 2 seconds. To obtain the regressors for the entire duration, the window was shifted by one sample, and the metric was recalculated again:

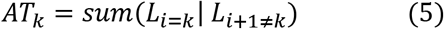

Similarly, the transition activity between microstate *m* and *l* is calculated as the number of switches between the two microstates before applying a sliding window of 2 seconds.

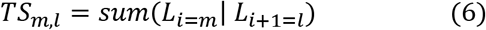

Finally, each regressor was convolved with double-gamma hemodynamic response function (Friston et al., 1998), standardized, and then down-sampled to the resting state repetition time (TR) of fMRI, resulting in EEG-ms-informed regressors to be included in BOLD signal analysis.

### GLM Analysis

Imaging analyses were carried out using the Analysis of Functional NeuroImages (AFNI) software (http://afni.nimh.nih.gov/afni/). The afni_proc.py command was employed to preprocess the data using the default parameters unless otherwise noted. The first three volumes were omitted from the analysis. The despike option was applied to avoid sudden jumps in the time courses of BOLD signals. RETROICOR (Glover, Li, & Ress, 2000) and respiration volume per time correction (Birn, Diamond, Smith, & Bandettini, 2006) were applied to remove cardiac- and respiration-induced noise in the BOLD signal. Slice-timing differences were adjusted by aligning to the first slice, and motion correction was applied by aligning all functional volumes to the first volume. EPI volumes were acquired using the 3dvolreg AFNI program with two-pass registration. The volume with the minimum outlier fraction of the short EPI dataset acquired immediately after the high-resolution anatomical (MPRAGE) brain image was used as the registration base. Linear warping was applied to Montreal Neurological Institute space and resampled to 2 mm3 voxels. Data were then smoothed using a Gaussian kernel of 6mm FWHM and then scaled to have a mean of 100 and range of [0, 200].

fMRI BOLD signal analysis was performed using the standard GLM approach with the AFNI 3dDeconvolve function (Cox, 1996). The design matrix included one EEG-ms regressor corresponding to one EEG-ms property (direct time, activity or transition) and a set of nuisance covariates: 1) low-frequency fluctuation from the signal time course (i.e., 3rd-order polynomial model); 2) 12 motion parameters (i.e., three-shift and three rotation parameters with their temporal derivatives); 3) three PCs of the ventricle signal from the signal time course; and 4) local white matter average signal (ANATICOR) (Jo, Saad, Simmons, Milbury, & Cox, 2010). GLM *β* coefficients were computed for each voxel, and then a one-sample *t*-test was applied to the entire sample of healthy subjects to extract EEG-ms templates. To control for potential false positives in BOLD signal (Eklund, Nichols, & Knutsson, 2016), 1) the non-Gaussian spatial autocorrelation function (ACF) was estimated for the dataset; 2) AFNI’s 3dClustSim was applied to the statistical map ((Cox, Chen, Glen, Reynolds, & Taylor, 2017); 3) a permutation test (n = 10000) was performed using the Smith procedure (Winkler, Ridgway, Webster, Smith, & Nichols, 2014), showing that an ACF-corrected cluster requires a minimum of 136 voxels to be deemed significant at *p* < 0.05—using an uncorrected voxel-wise threshold of *p* < 0.005. Moreover, the GLM model excluded TRs with severe motion (Root Mean Square (RMS) > 0.2) or with EEG artifact (i.e., if the TR was located within unusable intervals of EEG data). In addition to the previous steps, further exclusion criterion was applied to subjects’ fMRI datasets, given that the number of censored volumes. In details, we excluded any subject if the number of censored TRs was more than third of the total number of TRs. This was necessary to ensure that the GLM model had enough data to estimate beta coefficients.

## Results

### EEG-ms identification

We identified the four canonical EEG-ms, which replicate EEG-ms templates obtained in the literature (Michel & Koenig, 2018). The raw explained variance by the four EEG-ms was 73%, with 21%, 16%, 20%, and 16% for MS-A, MS-B, MS-C, and MS-D, respectively. **Figure 3** shows the extracted templates.

**Figure 3.**
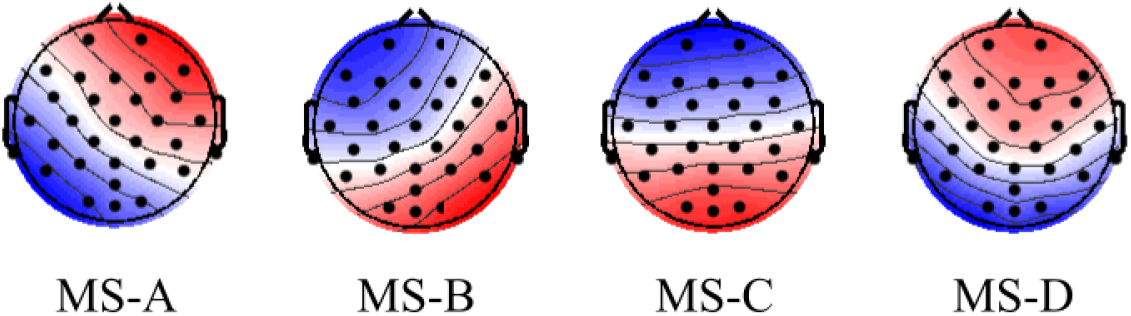
EEG-ms topographies obtained from the dataset.

Using Equation 3, we extracted the direct EEG-ms regressors. Due to the high correlation among those regressors (range= [-0.6, 0.95]), each regressor was used independently in the GLM model.

### Correlated fMRI maps of EEG-ms using direct time course regressors

### Correlated fMRI maps of EEG-ms activity

Using the activity regressors extracted based on Equation 5, we identified the correlated maps for each MS. As in the direct time course activity, each MS regressor was used independently in the GLM model. The results for the activity regressors are shown in **Figure 5** for MS-A, MS-B, MS-C, and MS-D.

To further characterize the spatial extent of EEG-ms correlated brain regions, we used the 7-network atlas provided by (Thomas Yeo et al., 2011). More specifically, the atlas provides the spatial distribution of somatomotor, visual, dorsal attention, ventral attention, limbic, frontoparietal, and default-mode networks. For each EEG-ms, we calculated the number of voxels situated within each network over the number of voxels within that network. **Fig. 6** shows the percentage of overlap for each EEG-ms across the seven networks and reveals significate overlap with somatomotor, visual, dorsal attention and ventral attention network.

### Correlated fMRI maps of EEG-ms transition regressors

EEG-ms lasts on average for ~50ms before switching to another microstate. Using the transition regressors (Equation 6), we ran a GLM using 16 transition regressors and identified the correlated maps (12 transitions among four microstates and four regressors of self-transitions). **Fig. 7, 8, 9, and 10** depict the correlated map with significant clusters for each EEG-ms (if present). It should be noted that *TS_A,D_* and all other within microstate-transitions (*TS_A,A_, TS_B,B_*, *TS_C,C_* and *TS_D,D_*) did not yield any significant clusters. All transitions have corresponding significant association except for *TS_A,D_*. Also, figures show different maps for symmetrical transitions (e.g., *TS_A,B_* and *TS_B,A_*).

## Discussion

We expanded efforts to understand the association among EEG-ms spatio-temporal dynamics and the BOLD fMRI signal and further understand the mechanism behind the presence of EEG-ms. In addition, we extracted the associated templates so they can be used further for studying and evaluating EEG-ms in the absence of fMRI. EEG-ms exhibit wide and stable topographical activation lasting for approximately 50 milliseconds. The large spatial distribution of EEG-ms, along with dynamic fluctuation among those EEG-ms topographies, offers valuable information about underlying neural processes in the brain, including cortical neuronal generators. EEG-ms dynamics are relativity rapid, with complex switching patterns among microstates. The common assumption about EEG-ms is that they represent coherent brain functions that raise to specific topographical representation. Herein, we inspected the association between simultaneously recorded EEG-ms and fMRI BOLD signal. We designed three sets of regressors to capture EEG-ms dynamics: (1) the direct time course of each microstate, which represents the spatial similarities between each EEG point and EEG-ms templates; (2) the switching activity from specific microstates to other microstates, which measures the extent of switching from one brain function to other states; and (3) the pairwise transitioning among microstates, which reveals more detailed information about dynamics among switching between brain functions. Overall, the result revealed a significant association between EEG-ms activity and the BOLD signal across several regions of the brain and might suggest that EEG-ms represent a sequence of activation/deactivation of brain functions rather than a specific brain function. We delineate the results for each type of regressors accordingly and offer the supporting discussion.

### EEG-ms direct time course regressors

Using the canonical EEG-ms direct time course regressors, nine significant clusters survived multiple-comparison correction: two clusters for MS-A, two clusters for MS-C, and five clusters for MS-D (**Fig. 4, Fig. S1 and Table 1**). The identified brain regions for MS-C (left middle frontal gyrus and cuneus) and MS-D (bilateral temporal gyri, left precentral and supramarginal gyri, and right cuneus) replicate findings for these MSs reported by (Britz et al., 2010). In addition, our results indicated that MS-A exhibits a negative association with bilateral lentiform gyri, while no brain regions were associated with MS-B. The number of voxels surviving our multiple-comparison correction was relatively smaller than reported activations by (Britz et al., 2010). This discrepancy might be due to our use of strict physiological and motion correction for resting-state analysis (RETROICOR and ventricle white matter regression). The previous results in (Britz et al., 2010) were extracted using [2-20] Hz filtering for EEG before applying EEG-ms analysis. Several works in the literature appeared to use [1-40] Hz filtering for EEG (Michel & Koenig, 2018). Thus, we investigated using regressors for [1-40] Hz range. The analysis revealed only two and four significant clusters for MS-C and MS-D, respectively (Supplementary: **Fig. S2** and **Fig. S3**). Although the spatial distribution of clusters was different from the [2-20] Hz case, no statistically significant differences were obtained when comparing the Beta maps for the two frequency configurations.

**Figure 4:**
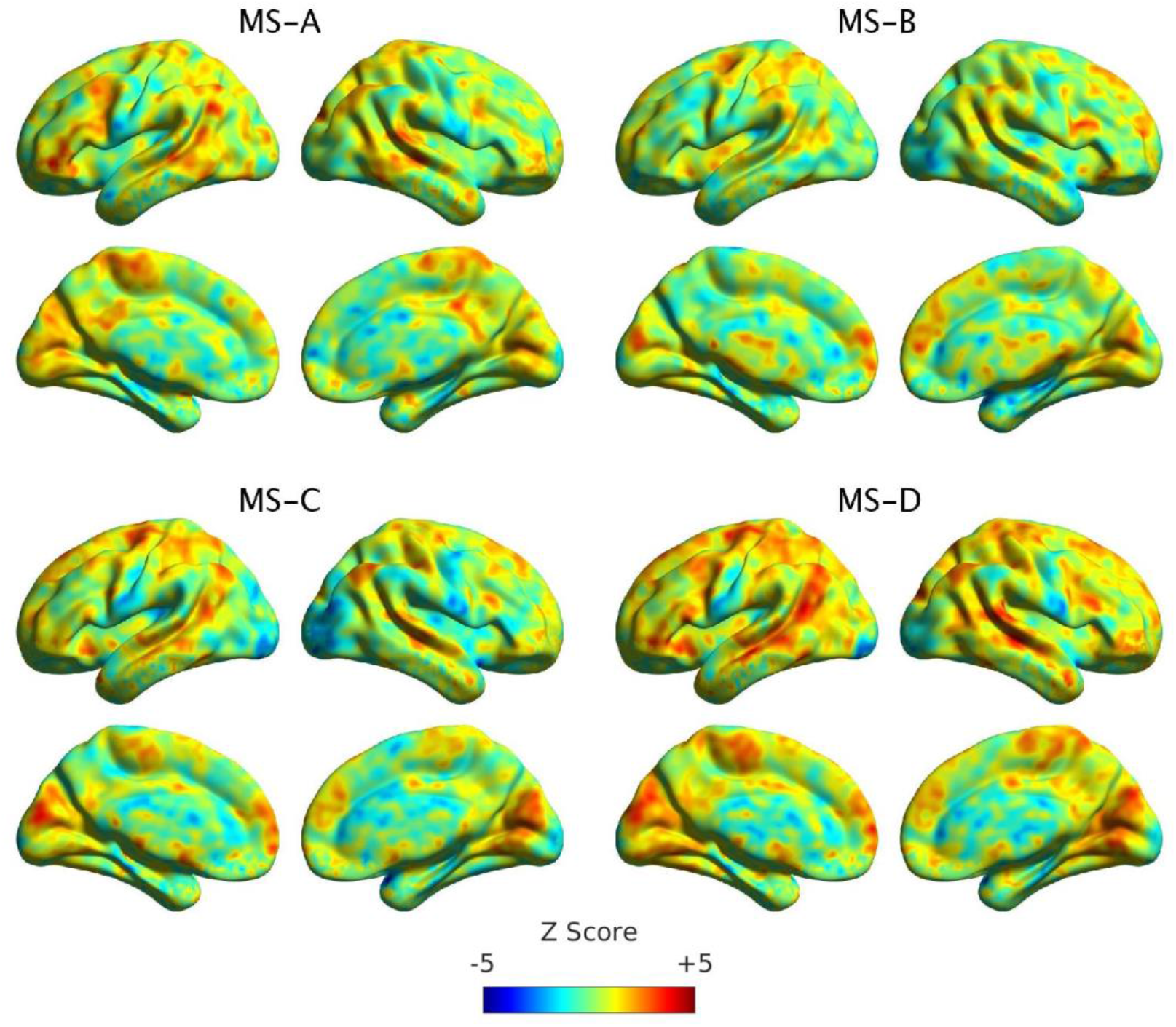
the un-thresholded maps of each EEG-ms direct time regressor. No significant clusters were found for MS-B (Please refer to the **Fig S1** in the supplement for thresholded clusters). Detailed information about each cluster and the corresponding brain regions are presented in **Table 1**.

**Figure 4.**
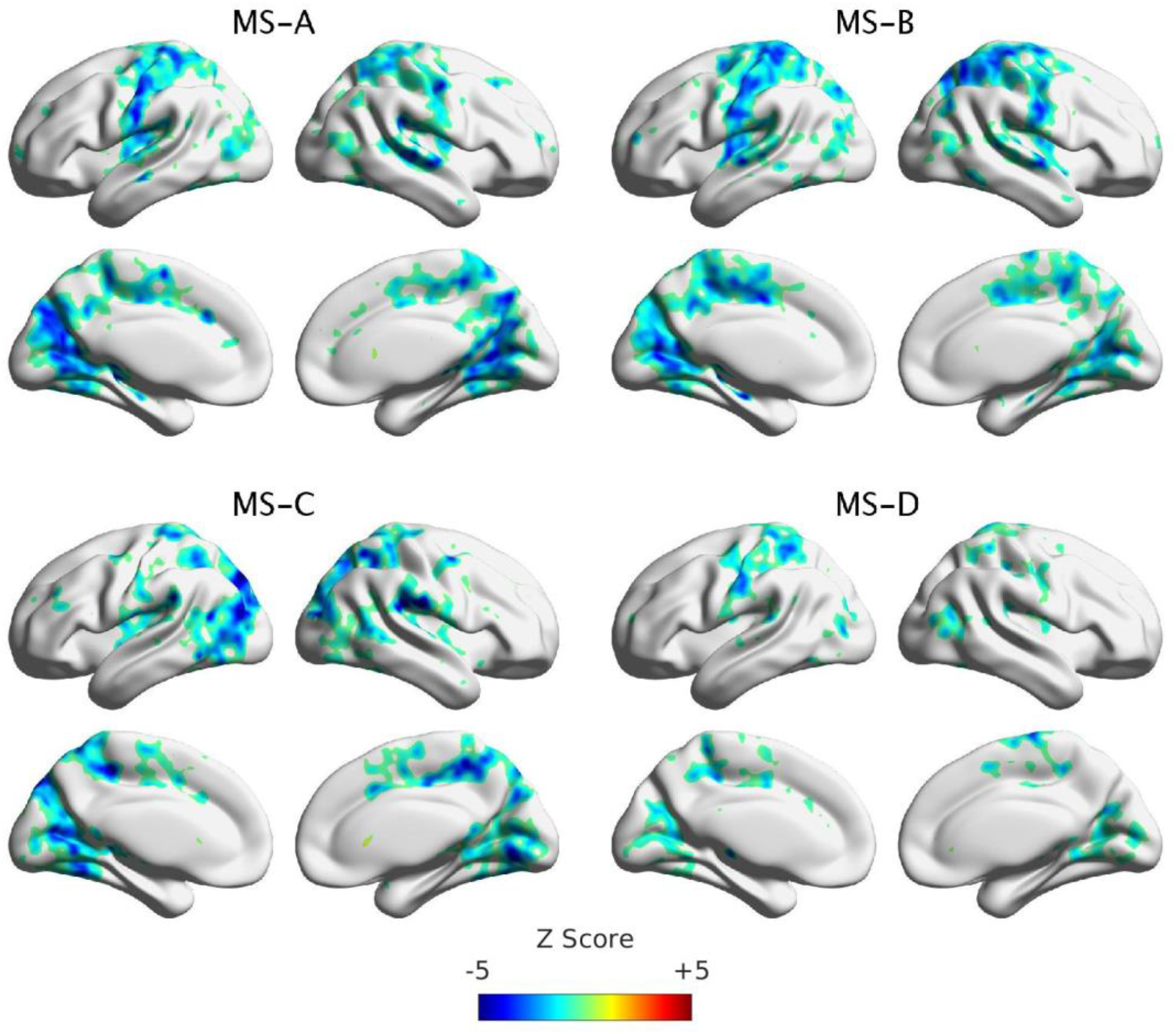
Significant clusters for MS-A, MS-B, MS-C, and MS-D using activity regressors. Clustering was performed at p<0.005 and corrected at p<0.05.

**Table 1.** Significant clusters using direct EEG-ms regressors. The coordinates are reported in MNI space.

| **MS** | **Cluster** | **Region** | **X** | **Y** | **Z** | **Voxel** |
| --- | --- | --- | --- | --- | --- | --- |
| A | C1 | Right Lentiform Nucleus | -26 | -2 | 2 | 231 |
| A | C2 | Left Lentiform Nucleus | 28 | -1 | -1 | 187 |
| C | C1 | Left Middle Frontal Gyrus | 28 | 11 | 64 | 206 |
| C | C2 | Left Cuneus | 2 | 85 | 20 | 176 |
| D | C1 | Right Cuneus | 0 | 82 | 27 | 470 |
| D | C2 | Left Supramarginal Gyrus | 50 | 58 | 26 | 361 |
| D | C3 | Left Precentral Gyrus | 25 | 16 | 64 | 244 |
| D | C4 | Left Middle Temporal Gyrus | 60 | 40 | -1 | 185 |
| D | C5 | Right Superior Temporal Gyrus | -56 | 38 | 13 | 165 |

### Activity regressors

The results from direct time course regressors suggested the need for assessing the activity of EEG-ms as regressors. Thus, we investigated the extent of association between EEG-ms activity per each microstate and the BOLD signal (**Fig. 5**). In other words, we tested whether EEG-ms switching is associated with a change in brain functions. The results revealed a mostly strong negative association between activity regressors and BOLD signals for all MSs. To identify the spatial maps of the correlated maps, we adopted the 7-network Atlas (Thomas Yeo et al., 2011) to compute the percentage of overlap with each network (**Fig. 6**). Results showed that both MS-A and MS-B considerably overlapped with the somatomotor network (57% and 63%, respectively). Moreover, MS-A overlapped with the visual network (40%), followed by dorsal attention and ventral attention. MS-B shares 36% with a dorsal attention network, followed by 36% for the visual network and 31% for the ventral attention network. MS-C expanded 45% of the visual network, followed by 38% for dorsal attention and 34% somatomotor networks. MS-D regions shared 28% with the somatomotor network, followed by 11% for dorsal attention and ventral attention networks. For the default mode network, there was a little overlap with 9%, 9%, 5%, and 1% for MS-A, MS-B, MS-C, and MS-D, respectively. MS-C shared a 7% overlap with the frontoparietal network, followed by 5% for MS-A and MS-B. It should also be noted that none of the MSs shared any overlap with the limbic network.

**Figure 5.**
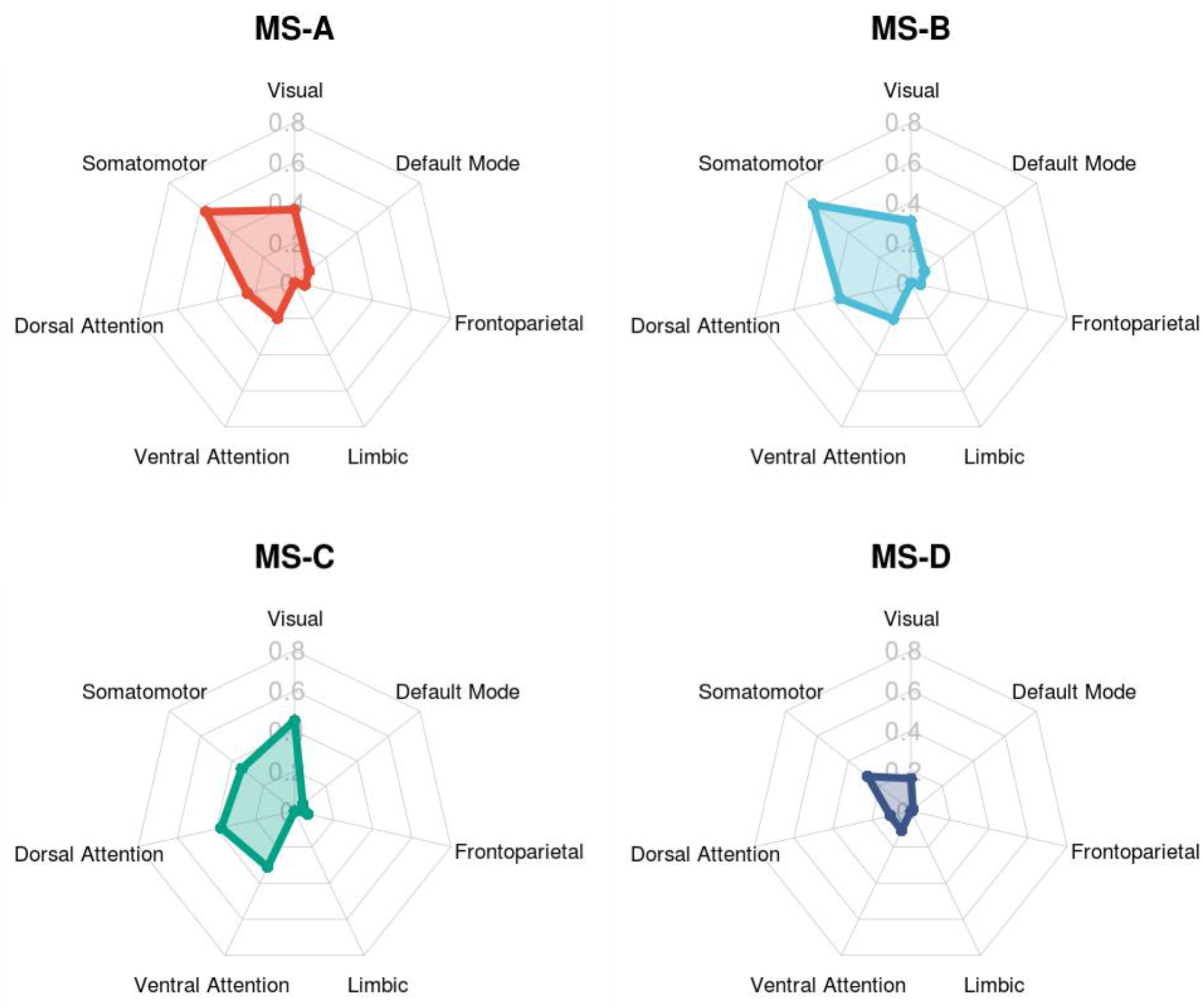
The percentage overlap between each EEG-ms correlated maps from activity regressors and the 7 brain networks identified by (Thomas Yeo et al., 2011).

**Figure 6.**
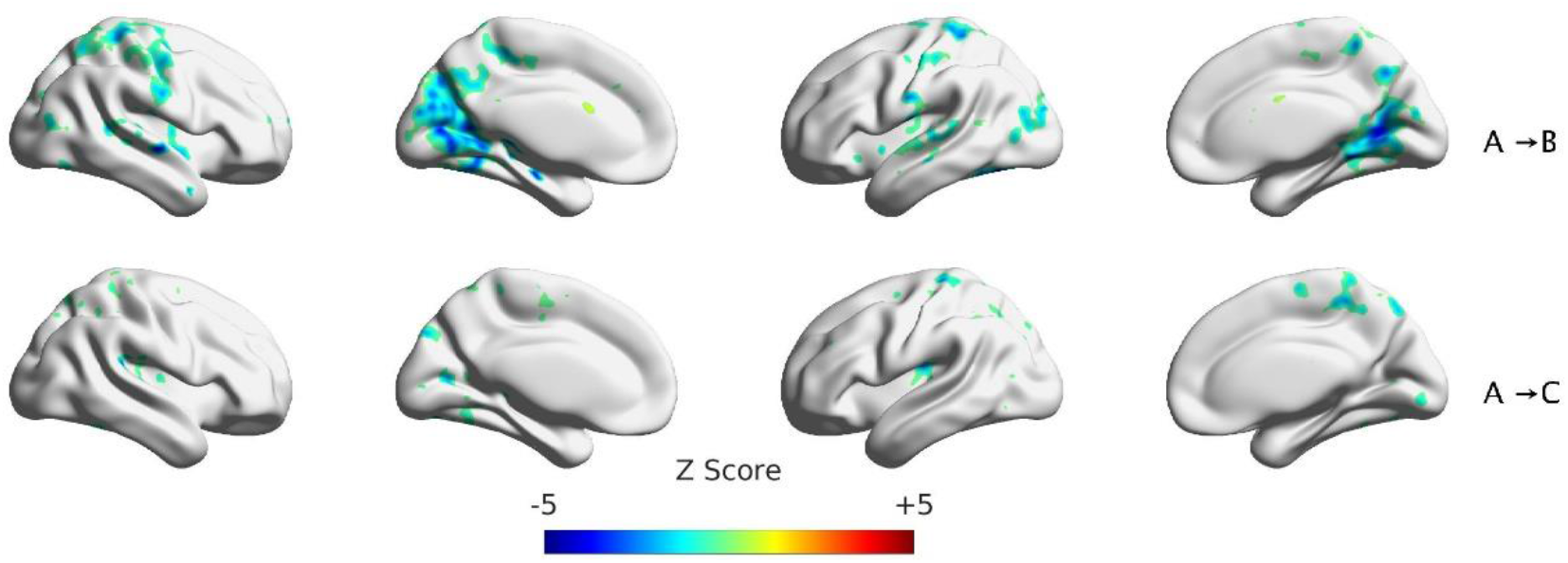
Significant clusters for transitions out of MS-A to other MSs. Clustering was performed at p<0.005 and corrected at p<0.05. It should be noted that there were no significant clusters for the A→D transition.

A similar negative association between EEG features and the BOLD signal has been found in the literature when using the alpha rhythm as a regressor (Goldman et al., 2002; Moosmann et al., 2003). More specifically, the alpha rhythm has been shown to be negatively correlated with the BOLD signal in the occipital brain regions. Additionally, EEG-ms classes have been shown to be predominantly impacted by the strength and spatial distribution of alpha rhythm (Milz et al., 2017). Also, the inter- and intra-subject variability of alpha rhythm along with another frequency modulates the EEG-ms metrics like occurrence, duration and coverage (Croce et al., 2020). There are consistently reported results about the role of alpha power in modulating the BOLD signal (Mantini et al., 2007; Scheeringa et al., 2012). Thus, this tri-relationship among BOLD, alpha rhythm and EEG-ms suggests that EEG-ms represent translational brain states that sum up to specific topographies. Also, this may explain the lack of spatial overlap between DMN and the spatial distribution of EEG-ms activity maps. The consistently reported evidence indicates that alpha rhythm oscillation exhibits inhibitory behavior in the task-related brain regions (Goldman et al., 2002; Milz, Pascual-Marqui, Lehmann, & Faber, 2016; G Pfurtscheller, 2003; Gert Pfurtscheller, Stancak Jr, & Neuper, 1996; Salenius et al., 1995; Scheeringa et al., 2012; Slatter, 1960). Thus, the lack of overlap with DMN, particularly when using activity regressors, may suggest that EEG-ms are associated with a sequence of deactivation of alpha rhythm and switching from task-negative to task-positive networks. This is also supported by the findings in (Milz et al., 2017; Pascual-Marqui et al., 2014). Also, this may explain the quasi-stability of EEG-ms classes and their coverage, where it is may be attributed to the lag in alpha rhythm changes when switching from task-negative to task-positive network reported in (de Pesters et al., 2016; Edwards et al., 2009; Potes, Brunner, Gunduz, Knight, & Schalk, 2014) and not as a result of invoking a specific brain function or network. Our interpretation of the dynamics among BOLD, EEG-ms, and alpha rhythm has an implication in interpreting previous EEG-ms literature, as has been noted to in (Milz et al., 2017). For instance, the prominence of MS-C during resting-state reported by (Milz, Faber, et al., 2016; Seitzman et al., 2017) may be explained as a strong deactivation of task-negative regions and invoking other brain functions. This is also supported by the fact that there is an increase in the alpha activity associated with MS-C reported in (Milz et al., 2017) and that alpha rhythm inhibition was associated with activating task-related brain regions. Other findings can be interpreted accordingly and provide an explanation for some inconsistencies like the differences among the spatial maps of MS-A and MS-B from (Britz et al., 2010) and the reported maps from (Milz, Faber, et al., 2016) by interpreting the maps as the extent of inhabitation rather than specific brain function utilization. To offer a complete discussion, we investigated using a [1-40] Hz configuration to extract the activity regressors. In this case, the analysis revealed a relatively similar activation pattern to the [2-20] Hz case (Supplementary: **Fig. S4**). However, comparing the Beta maps for the two frequency configurations revealed a change activation in the thalamic areas for MS A, B, and D with a higher activation for the upper frequencies (Supplementary: **Fig. S5**). Interestingly, a positive association between BOLD signal and alpha rhythm has been found for the thalamus as opposed to a negative association in the occipital brain regions (Goldman et al., 2002; Helmut Laufs et al., 2003; Moosmann et al., 2003; Pang & Robinson, 2018). In addition, a change in MS activation maps with upper frequency (>20 Hz) is consistent with reported results from (Schwab et al., 2015) for MS-B and MS-D, where authors reported a region-specific activation for subregions of the thalamus that were related to different MSs (although the authors did not report any thalamic activation for MS-A as in our case). To best of our knowledge, activity regressors have never been used in the literature to study the correlated maps between EEG-ms and BOLD signals. However, there might be shared information between the correlated maps of the temporal activity and sources of EEG-ms. Comparing the spatial distribution of activity regressors showed partial overlap with findings from an EEG-ms source-localization study (Custo et al., 2017). Our identified MS-A activity regions overlapped with the reported source localization of MS-A, and partially for MS-D. No overlap was found for MS-B and MS-C. However, it is difficult to draw conclusions due to methodological differences (Custo et al., 2017), such as extracting 7 MSs instead of 4 MSs and subtracting the mean of activation across the 7 MSs.

### Transition regressors

We also investigated the correlated maps of switching among pairs MSs by utilizing the transition among MSS as regressors. **Fig. 7, 8, 9**, and **10** unravel the correlated maps per each transition. Interestingly, the activity regressor did not show any significant brain regions for self-transition of EEG-ms (e.g., A→A), but a mostly negative association with BOLD signals for 11 of 12 transitions among different pairs of EEG-ms. It should be noted that the correlated maps of symmetrical transitions (e.g., A→B and B→A) have divergent patterns of activation and suggest that sequence of deactivation depends on the brain function that is being evoked. That is, if the symmetrical transitions have similar activation, one would assume that EEG-ms represent distinct brain functions. This is in line with our interpretation of EEG-ms using the interaction among BOLD, EEG-ms, and alpha rhythm. Furthermore, we studied the effect of using a [1-40] Hz frequency range to extract EEG-ms, which revealed similar activation for the transition regressors [2-20] Hz case and supports our interpretation of EEG-ms classes (See **Fig. S6, S7, S8** and **S9** in the Supplementary).

**Figure 7.**
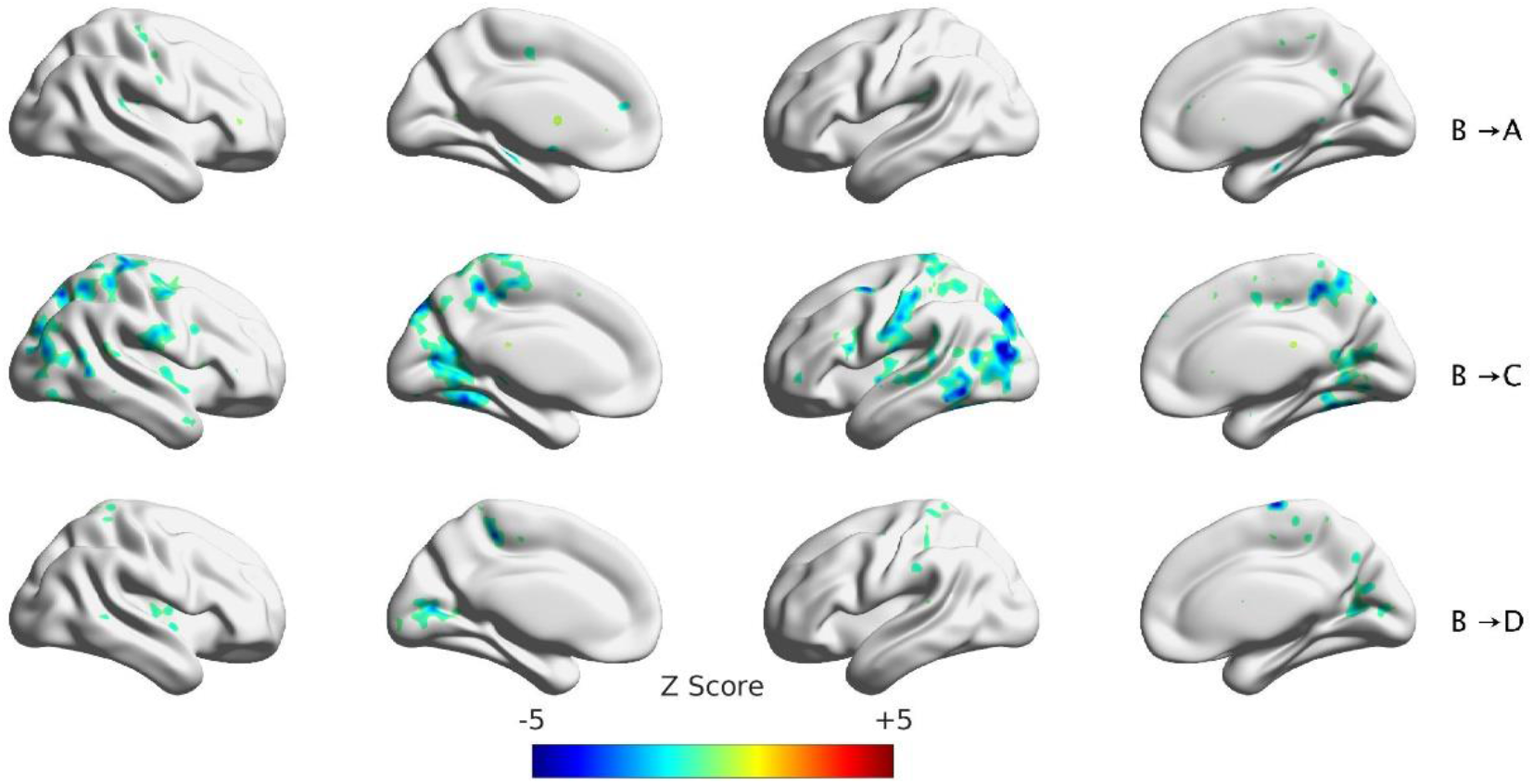
Significant clusters for transitions out of MS-B to other MSs. Clustering was performed at p<0.005 and corrected at p<0.05.

**Figure 8.**
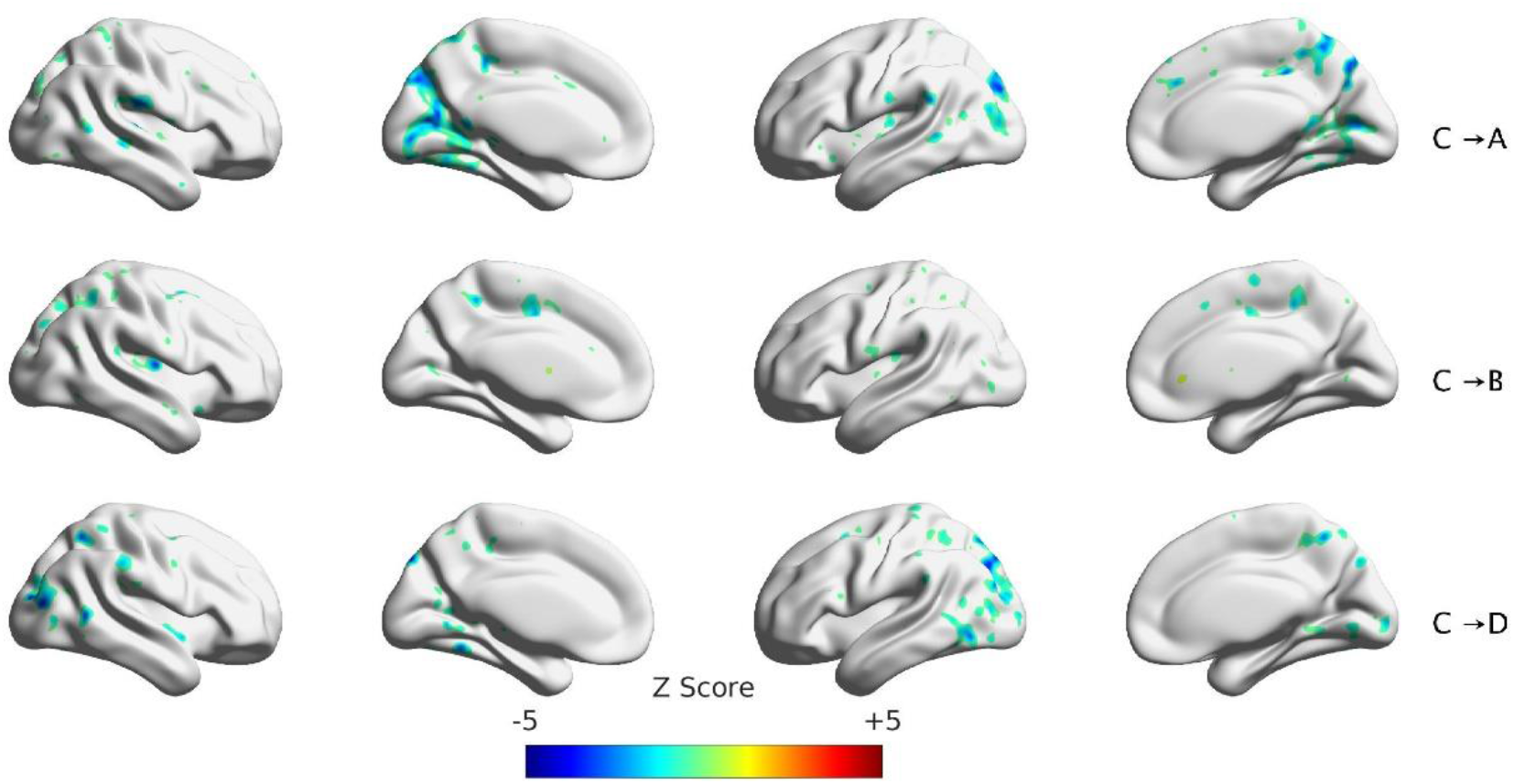
Significant clusters for transitions out of MS-C to other MSs. Clustering was performed at p<0.005 and corrected at p<0.05.

**Figure 9.**
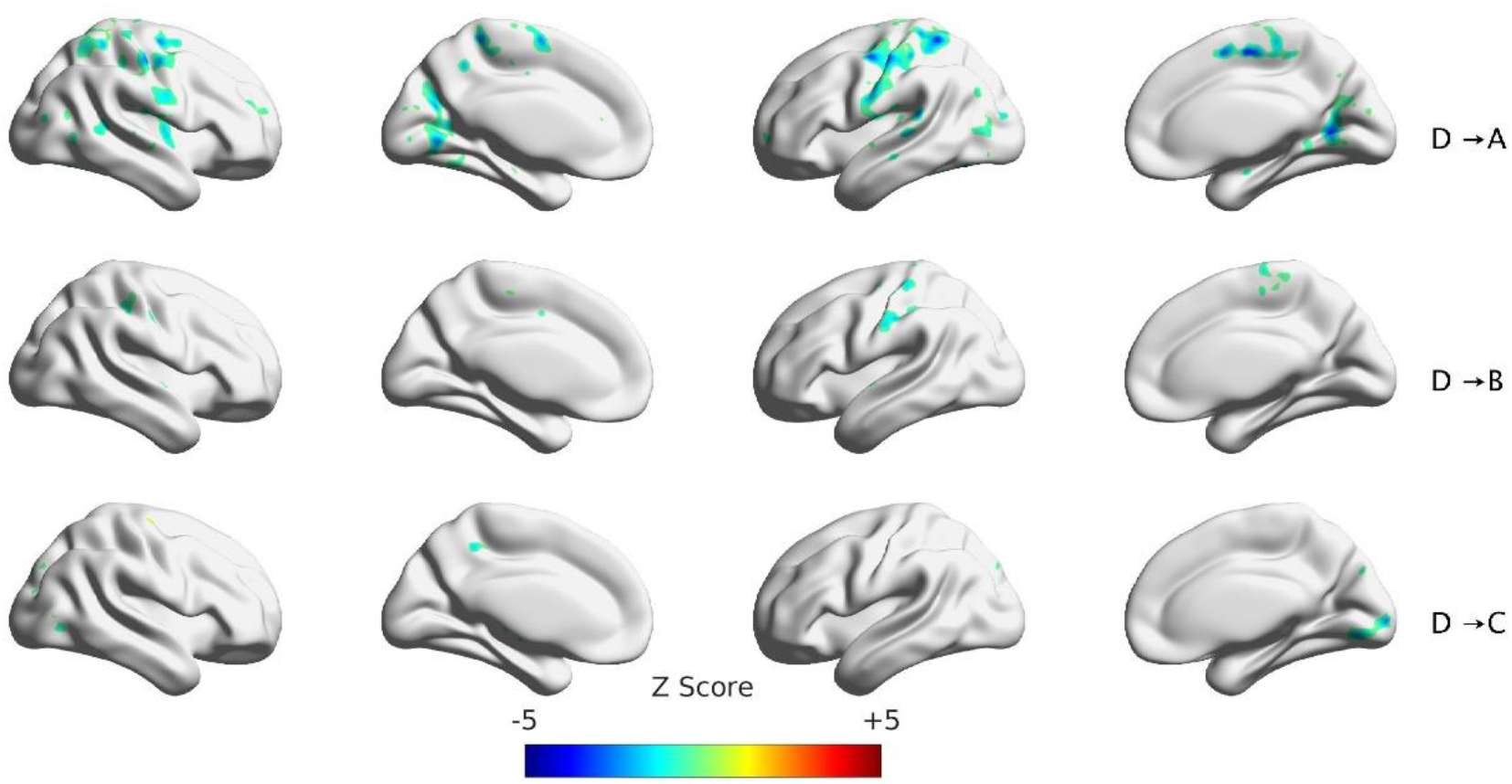
Significant clusters for transitions out of MS-D to other MSs. Clustering was performed at p<0.005 and corrected at p<0.05.

## Methodological Consideration

There are several limitations of the current work. First, the duration of the resting-state scan was 8 min. A longer duration might yield more robust estimations for the correlated maps and further avoid false positives. In addition, the analysis used a 32-channel EEG recording to conduct our analysis. High-density EEG recording might result in more robust results. Different number of TRs were used for different subjects due to censoring both EEG with bad interval points and fMRI with severe head motions. A comparison with other research was conducted based on the reported results. However, a more consistent approach should use the actual imaging data to compare the spatial distribution and the percentage of overlap. Finally, various works adopted some methodological procedures that differ from the canonical EEG-ms, which makes it difficult to compare results.

## Conclusion

EEG microstates (EEG-ms) have been used widely in the literature to characterize and to understand brain networks’ neuronal activities. In this work, we further expand the association between the dynamics of EEG-ms and the BOLD fMRI resting-state. First, we looked at the replication of previous studies that associated the EEG-ms time course and BOLD signal, where we found similar brain regions found in the previous studies, but with a smaller extent. Second, we investigated the association between EEG-ms temporal activity and BOLD, which have revealed a strong negative association between EEG-ms activity and BOLD signals focused on somatomotor, visual, dorsal attention, and ventral attention networks. A similar negative association was also found the pairwise transition regressors, but not for the self-transition. The results suggest that EEG-ms classes reflect switching from one brain functional state to another rather than being associated with specific brain functional state or RSN networks. Future studies should investigate in-depth the tri-relationship among BOLD, EEG-ms, and alpha rhythm.

## Supporting information

Supplement to EEG Microstates Mapping

## Availability of data

The data that support the findings and the data analysis scripts used in this study are available from the corresponding authors upon reasonable request.

## Acknowledgment

This work was supported by the Laureate Institute for Brain Research and the William K. Warren Foundation and in part by the P20 GM121312 award from the National Institute of General Medical Sciences, National Institutes of Health.

