## Supplement to EEG Microstates Mapping for "Canonical EEG Microstate Dynamic Properties and Their Associations with fMRI Signals at Resting Brain"

[*]The Tulsa 1000 Investigators include the following contributors:

Robin Aupperle, Ph.D., Jerzy Bodurka, Ph.D., Justin Feinstein, Ph.D., Sahib S. Khalsa, M.D., Ph.D., Rayus Kuplicki, Ph.D., Martin P. Paulus, M.D., Jonathan Savitz, Ph.D., Jennifer Stewart, Ph.D., Teresa A. Victor, Ph.D.

**EEG-ms Direct Time Courses Correlated Maps for 1-40 Hz filtering**

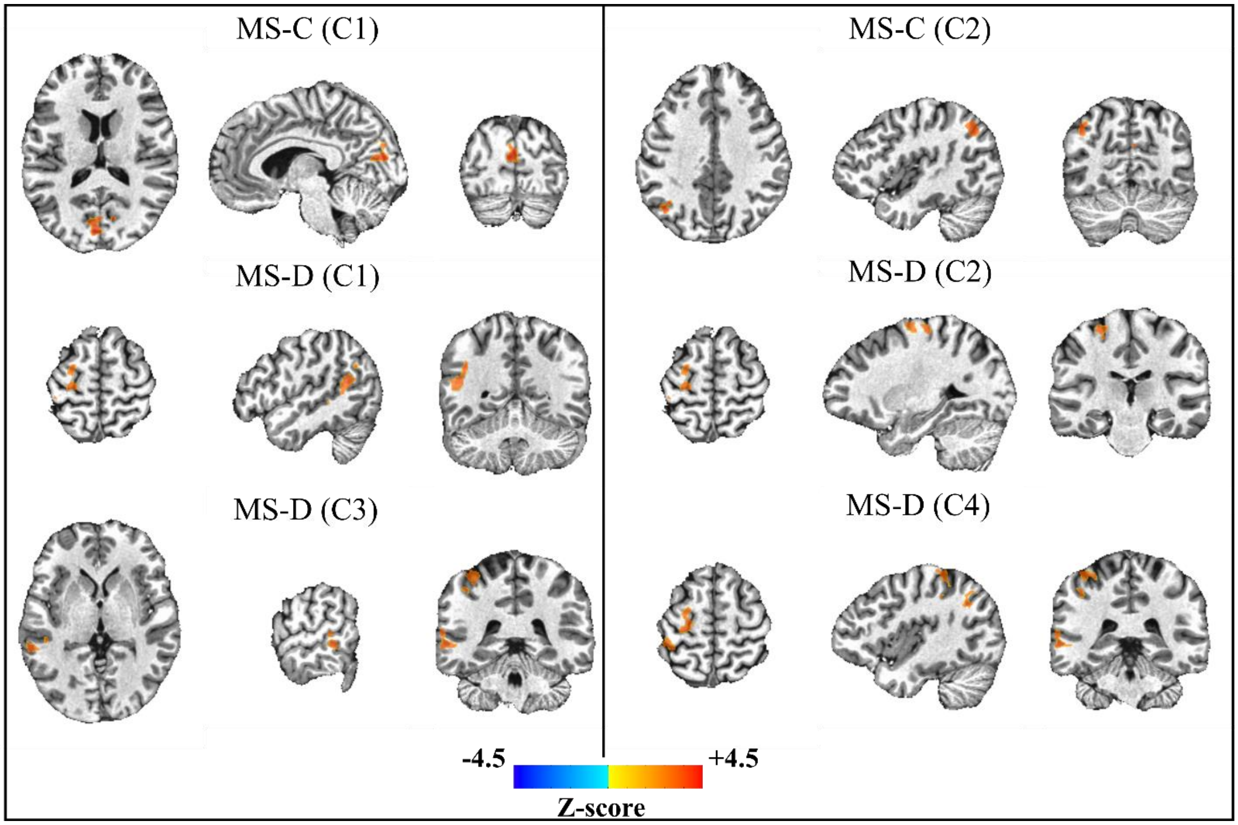


Figure S1: Significant clusters for MS-A, MS-C, and MS-D using fit regressors. Clustering was performed at p<0 .005 and corrected at p<0.05.

### EEG-ms Activity per microstate Correlated Maps for 1-40 Hz filtering

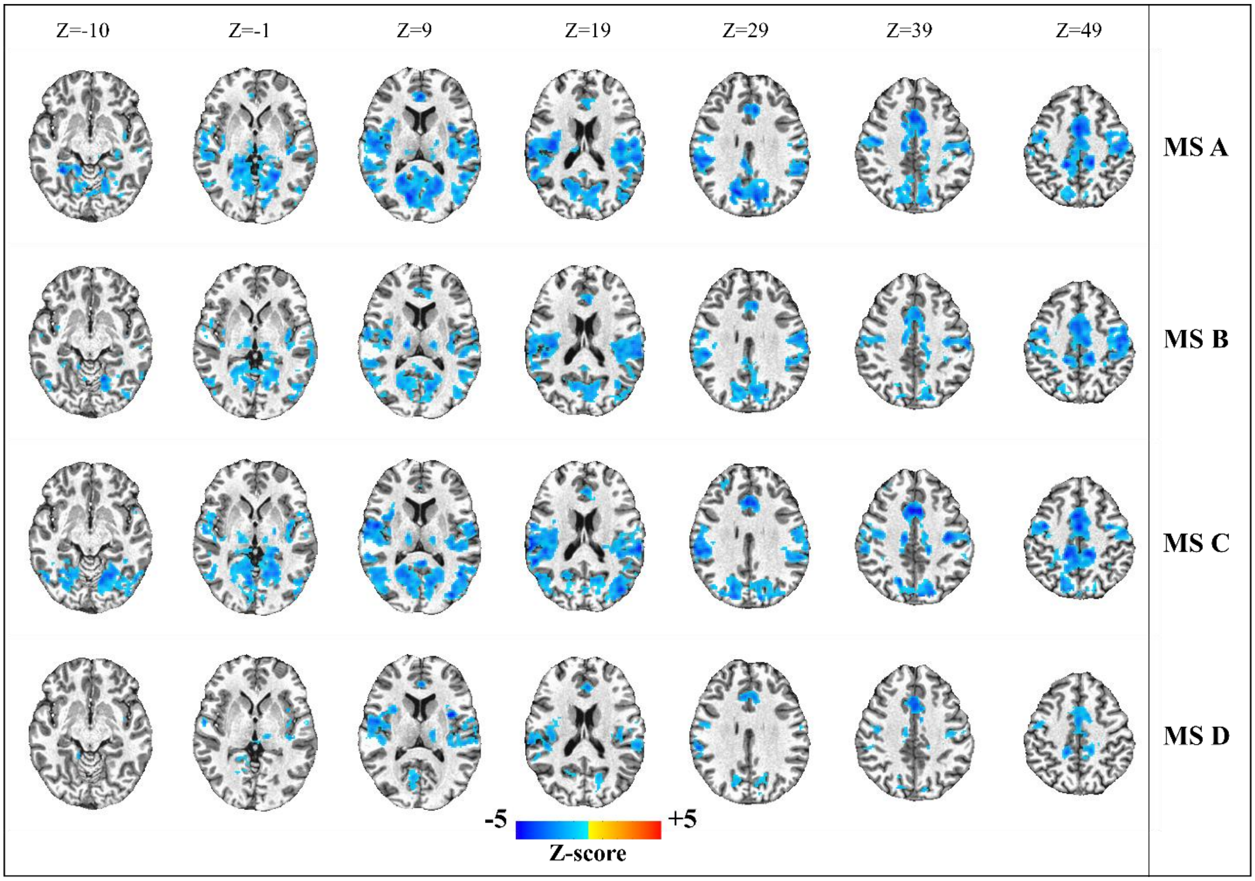


Figure S2: Significant clusters for MS-A, MS-B, MS-C, and MS-D using activity regressors. Clustering was performed at p<0 .005 and corrected at p<0.05.

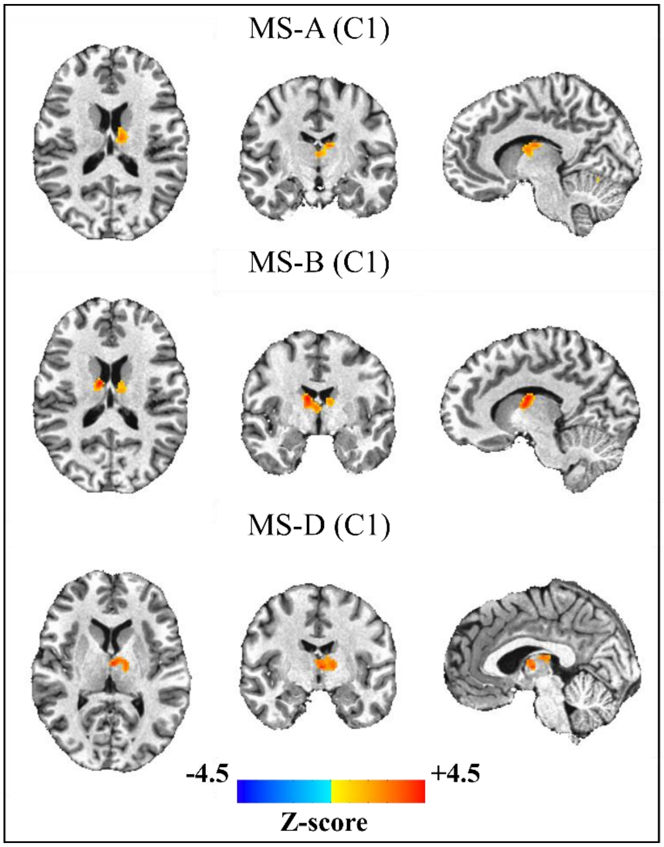


Figure S3: Significant clusters for paired t-test between [2-20] Hz filtering and [1-40] Hz. Clustering was performed at p<0 .005 and corrected at p<0.05.

### EEG-ms pair-wise transition per microstate Correlated Maps for 1-40 Hz filtering

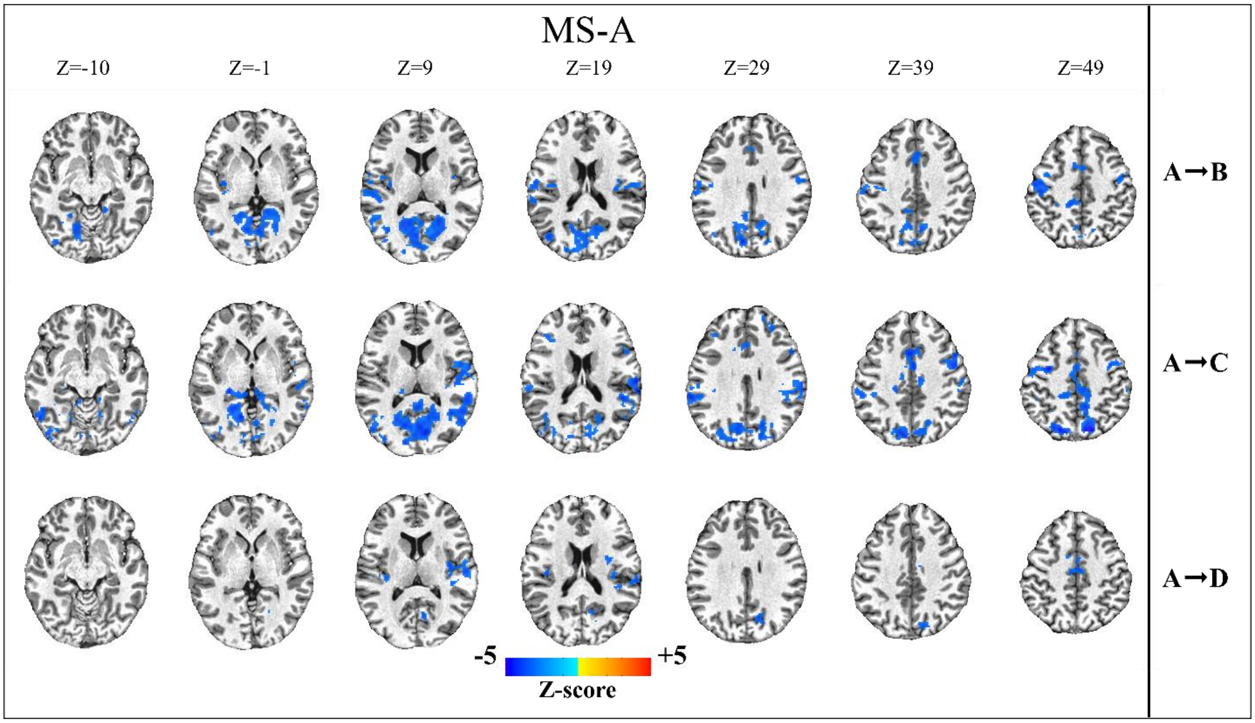


Figure S4: Significant clusters for transitions out of MS-A to other MSs. Clustering was performed at $p<0 .005$ and corrected at $p<0.05$.

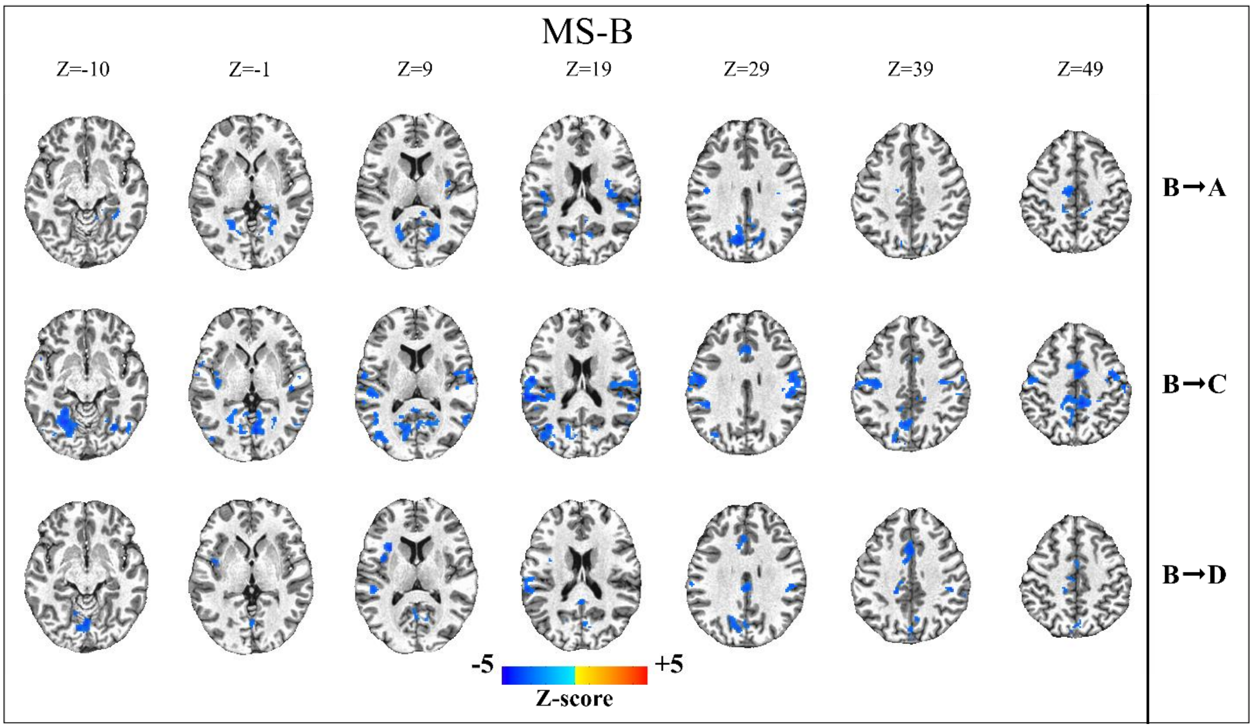


Figure S5: Significant clusters for transitions out of MS-B to other MSs. p<0 .005 and corrected at p<0.05.

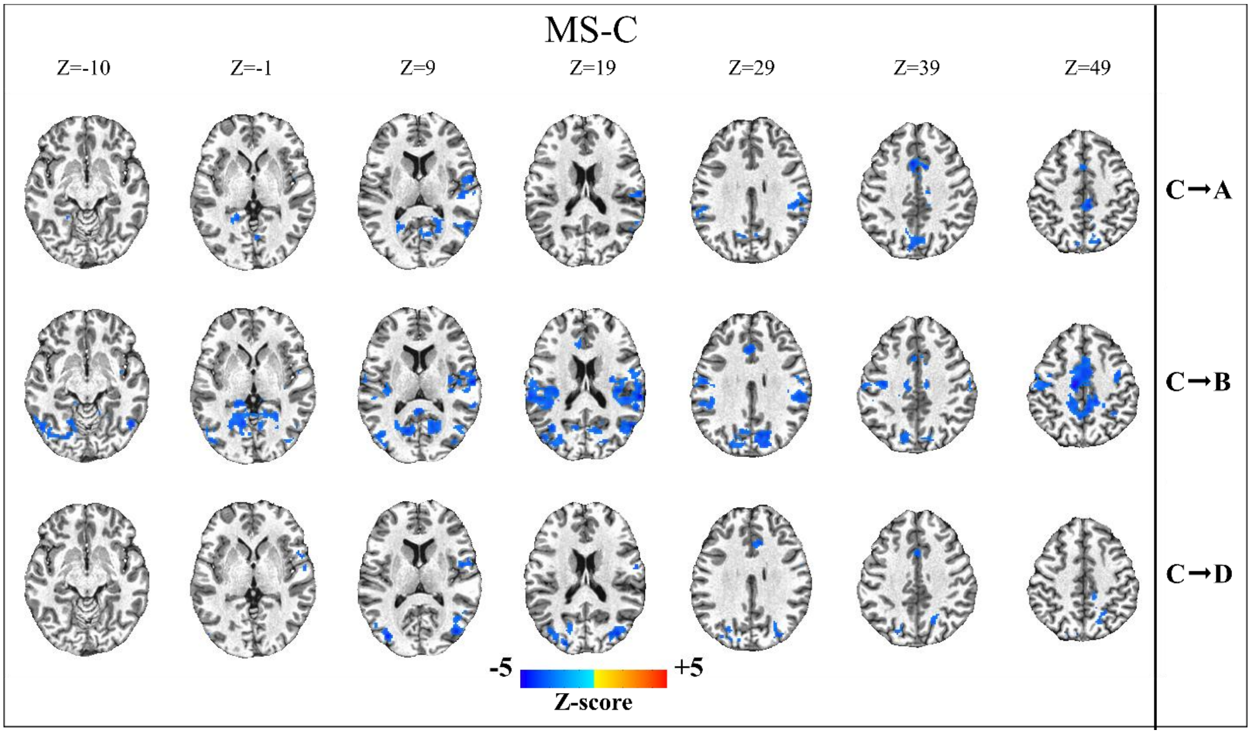


Figure S6: Significant clusters for transitions out of MS-C to other MSs. Clustering was performed at p<0 .005 and corrected at p<0.05.

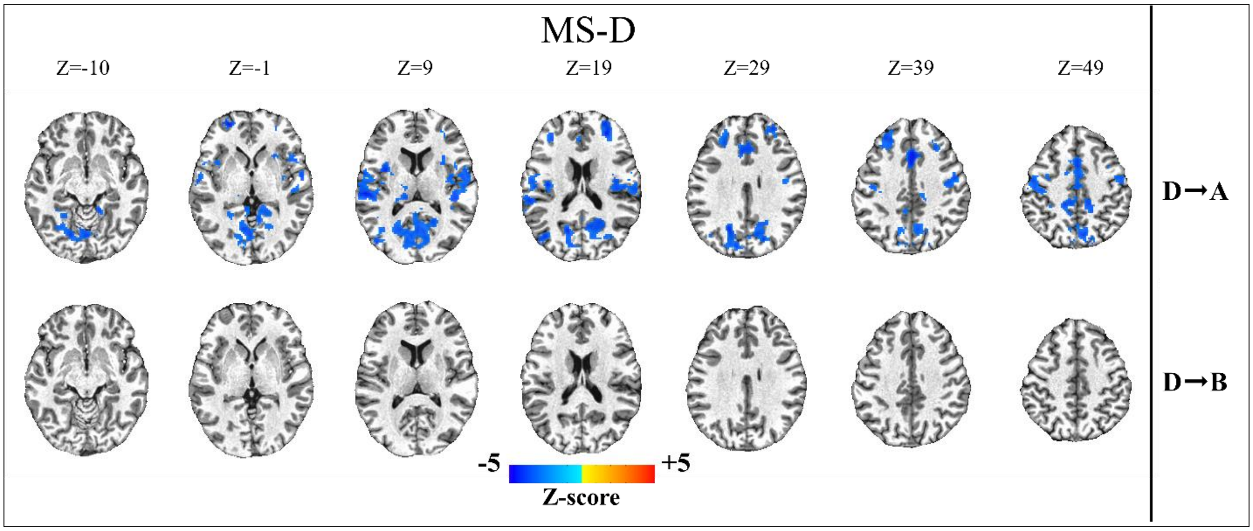


Figure S7: 10 Significant clusters for transitions out of MS-D to other MSs. Clustering was performed at p<0 .005 and corrected at p<0.05.
